# The rate of de novo CNVs in healthy controls

**DOI:** 10.1101/857797

**Authors:** Jacopo Barone, Mathew Smith, Kimberley Kendall, Michael J Owen, Michael C O’Donovan, George Kirov

## Abstract

**Background:** Copy number variation (CNV) is an important cause for human disease. Due to relatively high selection pressure operating against pathogenic CNVs, their rate is maintained in the population by *de novo* formation. The rates of *de novo* CNVs are increased in neurodevelopmental disorders. However only a few studies have been performed on relatively healthy individuals, making it problematic to calculate the magnitude of this increased rate.

**Methods:** The UK Biobank recruited about half a million randomly selected middle-aged members of the general population of the UK. We re-constructed family relationships from the genotypic data and identified 923 parent-offspring trios that passed out quality control filters. Potential *de novo* CNVs of >100 kb in size were identified and the log R ratios (LRR) and B allele frequency (BAF) traces of the trio members were visually inspected for those regions. We had no opportunity to validate CNVs with a laboratory method, but the traces appeared conclusive.

**Results and Discussion:** We identified 10 CNVs >100kb in size, a rate of 1.1%. These rates are very similar to those in previous large studies. Using previous large studies, we provide overall rates among 4844 trios for different size ranges that are expected in relatively healthy populations. These rates can be used for comparison in studies on disease populations.

## Introduction

Copy number variants (CNVs) are chromosomal deletions and duplications that range in size from kilobases to megabases of DNA sequence (Kirov, 2015). Research has implicated the role of large, rare CNVs as risk factors for schizophrenia, autism ASD, developmental delay and other neurodevelopmental disorders (Rees et al, 2014).

*De novo* mutations found in probands with a particular disorder can implicate them directly, as long as the rate of such *de novo* mutations in probands is higher than that in the general population. It is usually not possible for researchers to find sufficient numbers of control (unaffected) trios, in order to compare them with their disorder of interest. Most projects have instead to rely on established mutation rates in order to draw conclusions. The number of unaffected trios reported in the literature for *de novo* CNV analysis is relatively small (Xu et al, 2008; Malhotra et al, 2011; Kirov et al, 2012; Georgieva et al, 2014) and therefore there is a need to establish more robust estimates. The UK Biobank presents such an opportunity. It recruited half a million middle aged people from the general population of the UK and genotyped them with Affymetrix arrays that allow CNV analysis (Owen et al, 2018). Recruiting families was not the object of this initiative, but with a high proportion of people from certain areas being recruited, it was inevitable that some full trios had been included.

## Methods

### Participants & Genotyping

The UK Biobank recruited people from the general population of the UK, using National Health Service patient registers, with no exclusion criteria (Allen et al, 2012). Participants have consented to provide personal and health information, urine, saliva and blood samples, and to have their DNA tested. Samples were genotyped at Affymetrix Research Services Laboratory, Santa Clara, CA. Approximately 50,000 samples were genotyped on the UK BiLEVE Array (807,411 probes), with the rest on the UK Biobank Axiom Array (820,967 probes). There is 95% common content between the two arrays. CNV calling by our group has been described in previous work (Kendall et al, 2017, Crawford et al, 2019). Approval for our study on CNVs was obtained from the UK Biobank under project 14421: “Identifying the spectrum of biomedical traits in adults with pathogenic copy number variants (CNVs)”.

### Identification of trios

We used the released kinship coefficients and identity by descent (IBS0) data from the UK Biobank, following these rules: a trio had to consist of one person (proband) who is related in a first-degree relationship (kinship coefficient of ∼0.25) to two other people of different sex, who are >15 years older than the proband. The proband had to have IBS0 level close to zero with these two people (i.e. a parent cannot have alleles AA, while the child alleles BB at a locus). The two potential parents had to be unrelated. All three members of the trio had to pass our standard CNV QC criteria (genotyping call rate>0.96, number of CNV call per person <31, waviness factor >-0.03 & <0.03 and LRR-SD <0.35. We excluded probands with a diagnosis of developmental delay, autism, schizophrenia or psychosis, as we wanted to establish the de novo rate among individuals unaffected with early neurodevelopmental disorders. We identified 923 trios that satisfied all criteria.

### CNV calling

Our methods have been detailed before (Kendall et al, 2017; Crawford et al, 2018). Individual CNVs were excluded if they were covered by <15 probes, were <100kb in length, had a confidence score <10, a density coverage of <1 probe per 20 kb, low copy repeat (LCR) cover of >50% and a frequency > 1% in the sample as a whole. All potential *de novo* CNVs were assessed via visual inspection of the BAF and LRR plots of the three members of the trio. We could not validate results with another laboratory method but the BAF and LRR plots appear definitive, to leave us reasonably confident that they are real (Supplementary Material).

## Results

We confirmed 10 *de novo* CNVs >100kb in size: seven deletions and three duplications (Table 1). Of those, three are in known pathogenic loci.

**Table 1.**
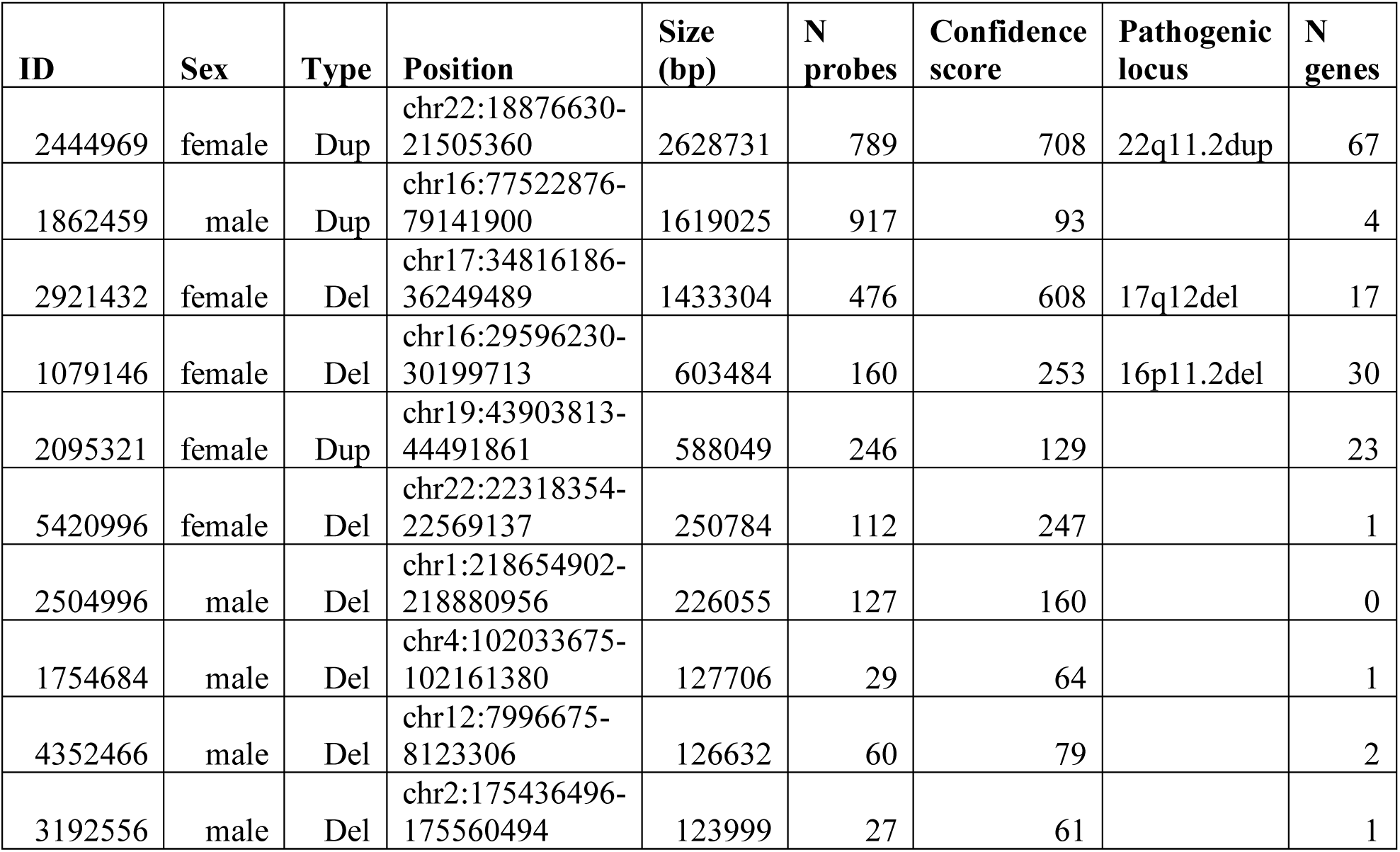
List of the 10 *de novo* CNVs, ordered by size.

| ID | Sex | Type | Position | Size (bp) | N probes | Confidence score | Pathogenic locus | N genes |
| --- | --- | --- | --- | --- | --- | --- | --- | --- |
| 2444969 | female | Dup | chr22:18876630-21505360 | 2628731 | 789 | 708 | 22q11.2dup | 67 |
| 1862459 | male | Dup | chr16:77522876-79141900 | 1619025 | 917 | 93 |  | 4 |
| 2921432 | female | Del | chr17:34816186-36249489 | 1433304 | 476 | 608 | 17q12del | 17 |
| 1079146 | female | Del | chr16:29596230-30199713 | 603484 | 160 | 253 | 16p11.2del | 30 |
| 2095321 | female | Dup | chr19:43903813-44491861 | 588049 | 246 | 129 |  | 23 |
| 5420996 | female | Del | chr22:22318354-22569137 | 250784 | 112 | 247 |  | 1 |
| 2504996 | male | Del | chr1:218654902-218880956 | 226055 | 127 | 160 |  | 0 |
| 1754684 | male | Del | chr4:102033675-102161380 | 127706 | 29 | 64 |  | 1 |
| 4352466 | male | Del | chr12:7996675-8123306 | 126632 | 60 | 79 |  | 2 |
| 3192556 | male | Del | chr2:175436496-175560494 | 123999 | 27 | 61 |  | 1 |

The rates of *de novo* CNVs in the UK Biobank were similar to those reported in previous studies (Table 2). We only included the largest available studies, to avoid potential publication bias and simplify the presentation.

**Table 2.** Comparison of rates of CNVs with previous studies on *de novo* CNV. The results from the current study are shown in bold.

| Study | Array | N | >100kb | >200kb | >500kb | >1Mb |
| --- | --- | --- | --- | --- | --- | --- |
| Sanders et al, (2011) | Illumina 1M | 872 unaffected siblings | N=11<br>1.3% | N=7<br>0.8% | N=7<br>0.8% | N=4<br>0.5% |
| DeCODE Controls (Kirov et al, 2012) | Illumina 317 (59.2%)<br>Illumina 370 (32.9%)<br>Illumina 1M (7.9%) | 2623 controls | N=46<br>1.8% | N=33<br>1.3% | N=20<br>0.8% | N=9<br>0.3% |
| Malhotra et al, (2011) | Nimblegen HD2 | 426 controls | N=1<br>0.2% | N=1<br>0.2% | N=1<br>0.2% | N=1<br>0.2% |
| Totals in previous studies |  | 3921 trios | N=58<br>1.5% | N=41<br>1.0% | N=28<br>0.7% | N=14<br>0.4% |
| **UK Biobank** | **Affymetrix Axiom** | **923 controls** | **N=10**<br>**1.1%** | **N=7**<br>**0.8%** | **N=5**<br>**0.6%** | **N=3**<br>**0.3%** |
| Overall rates |  | 4844 controls | N=68<br>1.4% | N=48<br>1.0% | N=33<br>0.7% | N=17<br>0.4% |

## Discussion

The goal of the present project was to obtain information on *de novo* CNV formation in a new large control sample, in order to establish a more reliable overall rate. Although previous large studies employed control groups from different settings and used different arrays (Malhotra et al, 2011; Sanders et al, 2011; Kirov et al, 2012), the UK Biobank data produced similar overall rates for different size cut-offs. Although a cut-off of >200kb in size is likely to produce more robust findings, the data show that for >100kb size the rates are also very similar, indicating that calls >100kb are also reliable.

The rates of 1.4% for CNVs >100kb, 1% for CNVs >200kb and 0.4% for those >1Mb only apply to relatively healthy adults and young people. This rate is bound to be higher among children and new-borns, as a proportion of those will develop ASD, ID, schizophrenia and other disorders that will make unlikely to be recruited as controls in genetic studies. We have shown that this effect on the overall rates is small for most CNVs that have incomplete penetrance, leading to similar rates in healthy controls and new-borns (Kirov et al, 2014). The effect differs for individual loci, depending on their penetrance. An almost fully penetrant CNV, such as 22q11.2 deletion, will only rarely be observed among controls from the general population (e.g. only 10 carriers in the UK Biobank, instead of the expected >100 among a populations of new-borns). However, most *de novo* CNVs in the general population are of incomplete penetrance, so as a group, the reported rates in this study (1.4% for CNVs >100kb) should be close to the expected rates in new-borns. Therefore, even for studies examining the rate of *de novo* CNVs among adults or young people with a specific disease, the control rates found in the current study should provide a close comparison.

## Supporting information

Supplementary Maaterial

## Acknowledgements

This research has been conducted using the UK Biobank Resource under Application Number 14421.

## Conflict of interest statement

The work at Cardiff University was funded by the Medical Research Council (MRC) Centre Grant (MR/L010305/1) and Program Grant (G0800509).

