## Supplementary Maaterial for "The rate of de novo CNVs in healthy controls"

Chromosome 16, ID 3022929, position 27,596,143 to 32,199,809

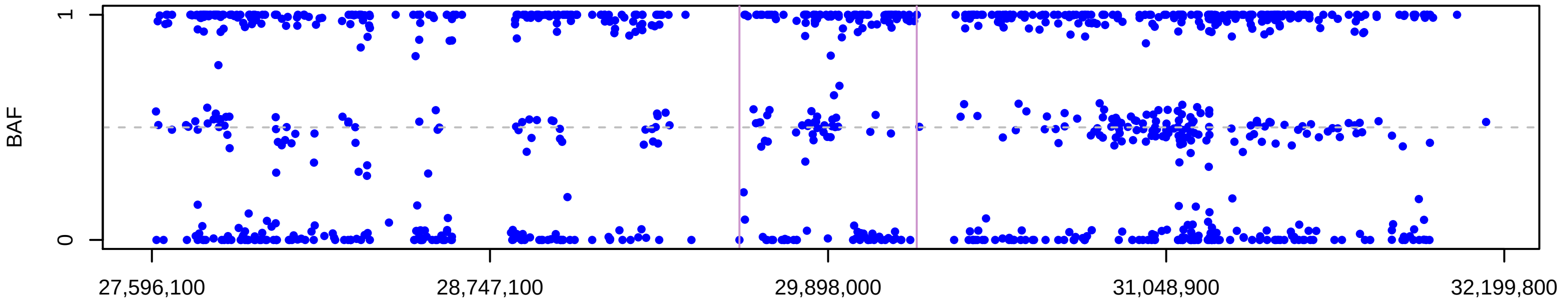


Chromosome 16, ID 3022929, position 27,596,143 to 32,199,809

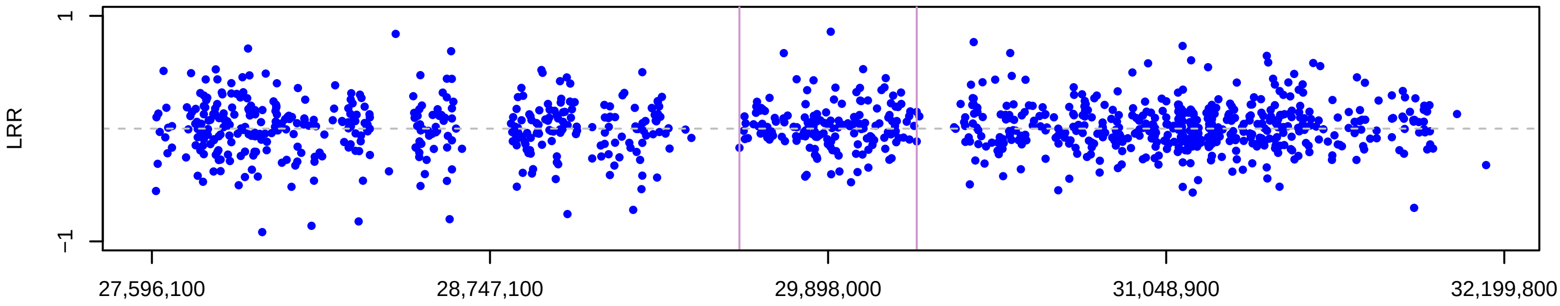


Chromosome 16, ID 3605380, position 27,596,143 to 32,199,809

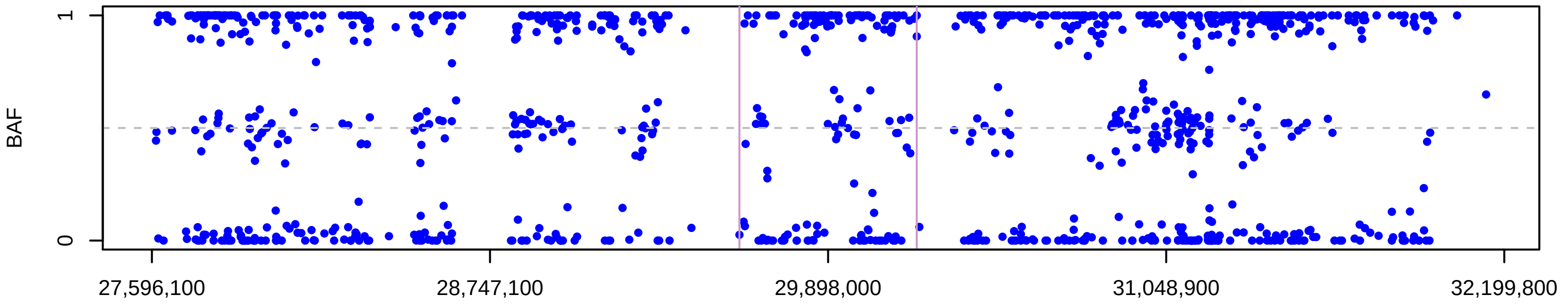


Chromosome 16, ID 3605380, position 27,596,143 to 32,199,809

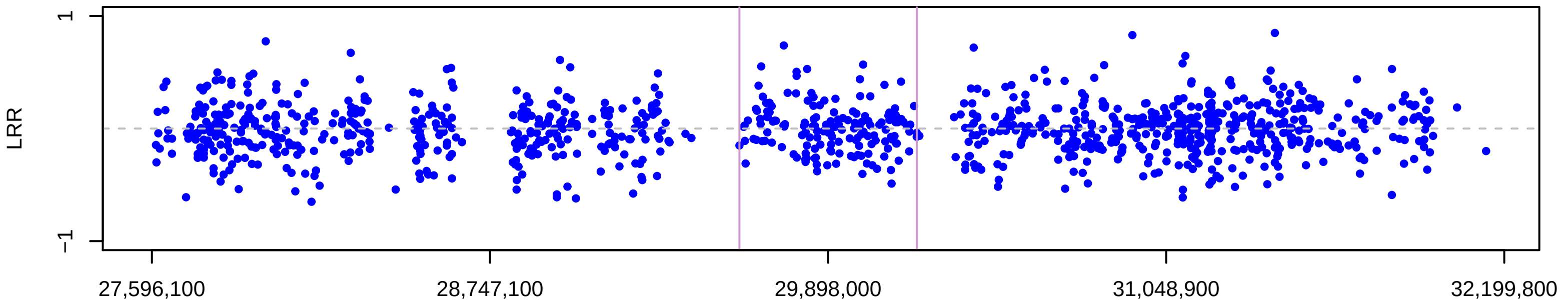


Chromosome 16, ID 1079146, position 27,596,143 to 32,199,809

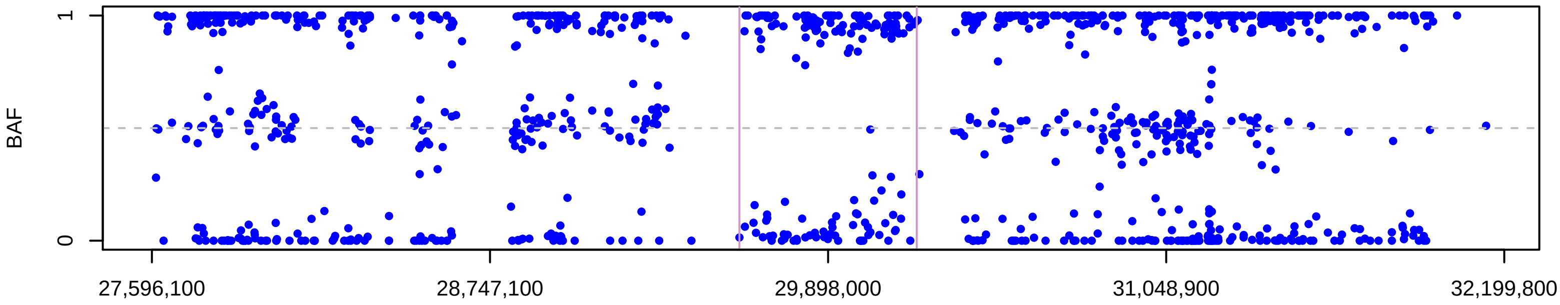


Chromosome 16, ID 1079146, position 27,596,143 to 32,199,809

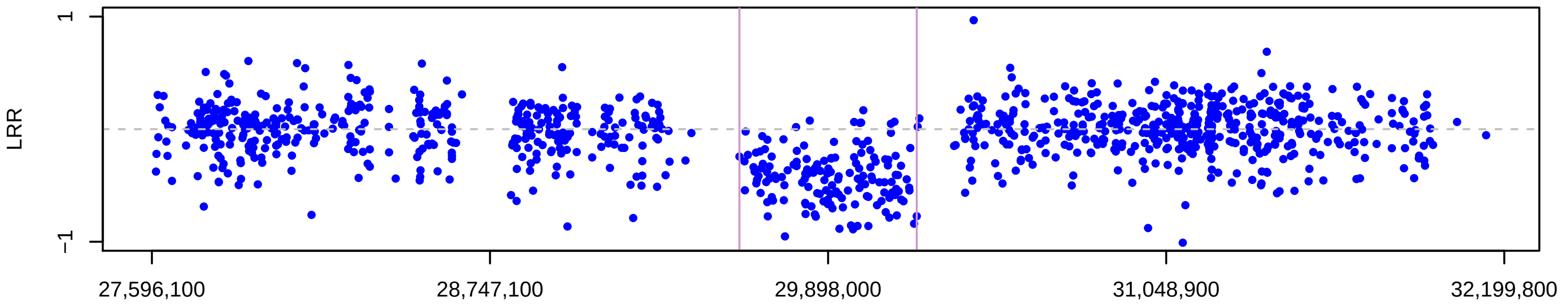

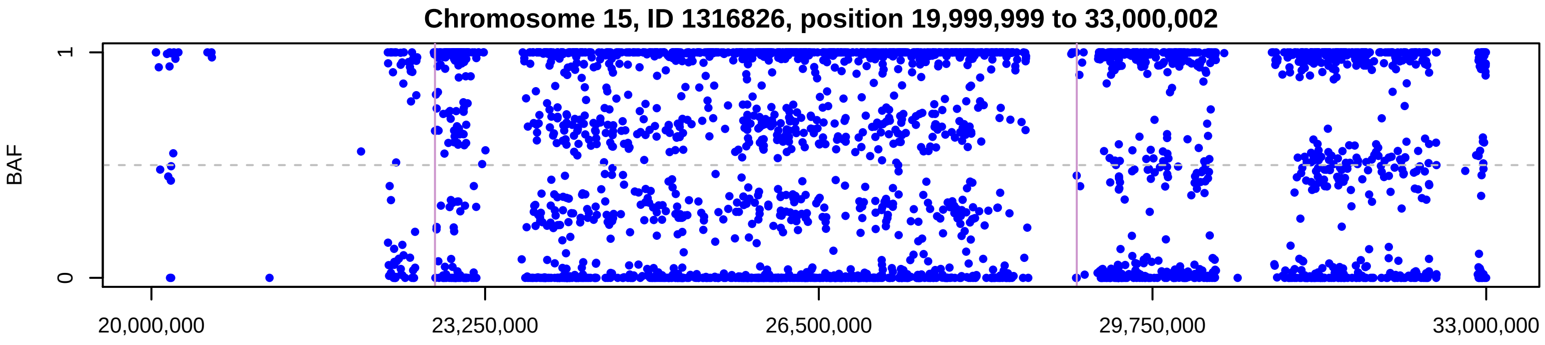


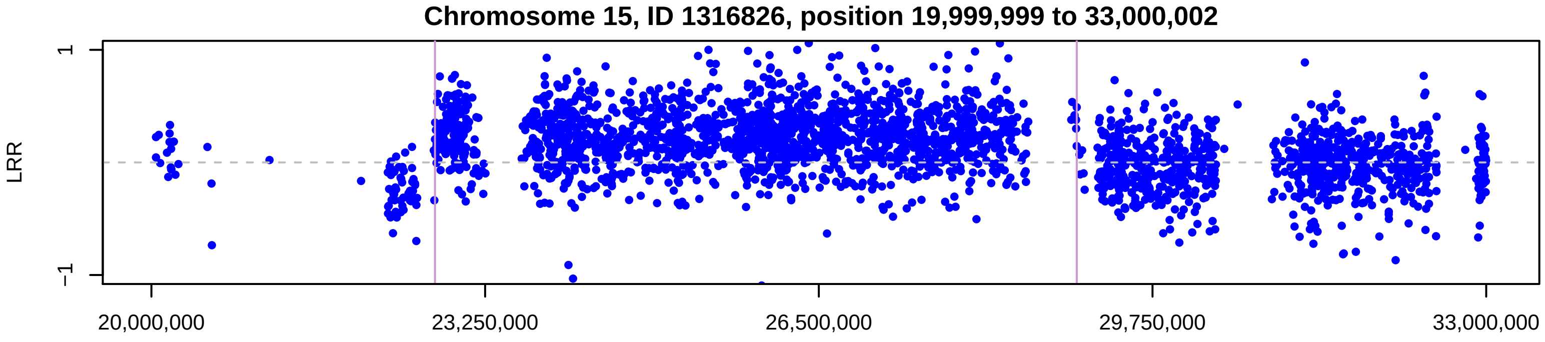


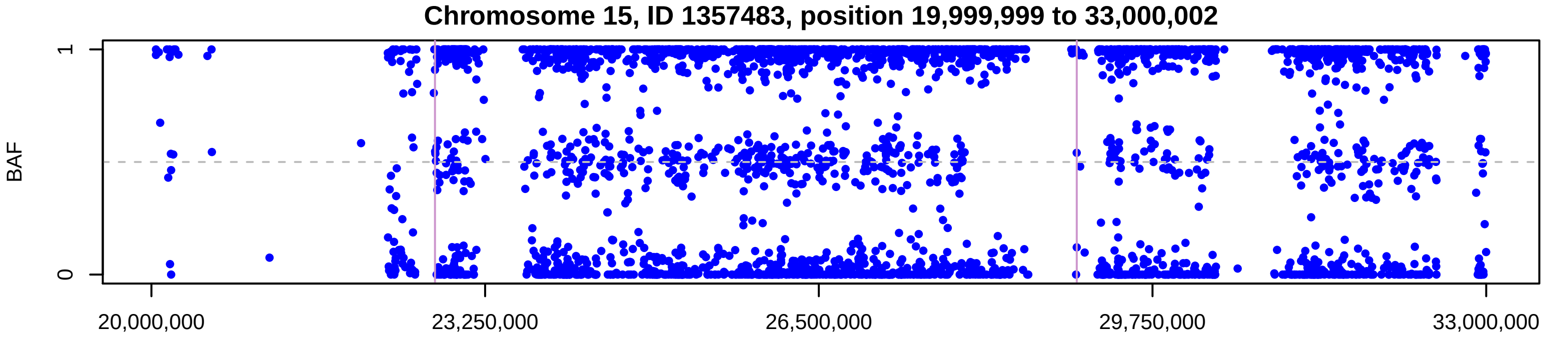


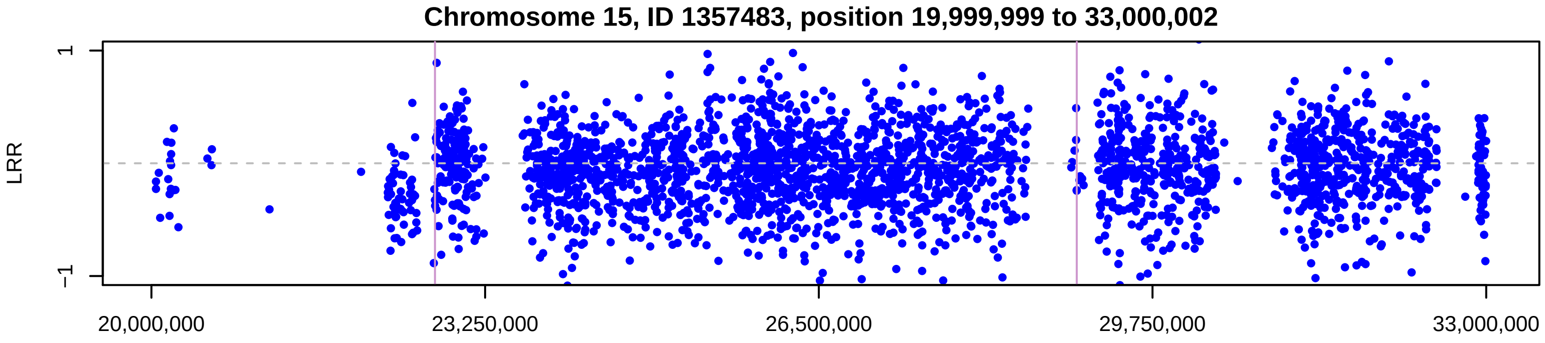


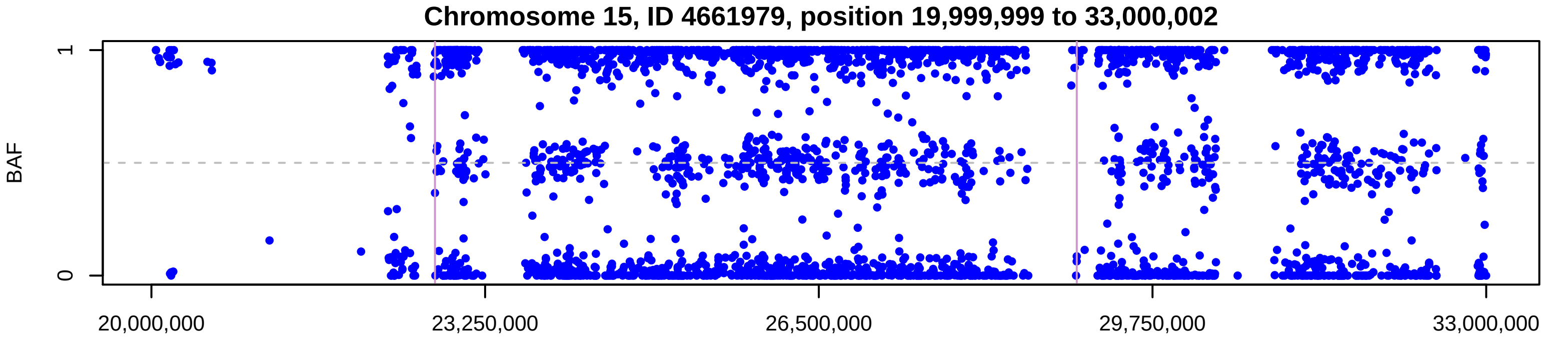


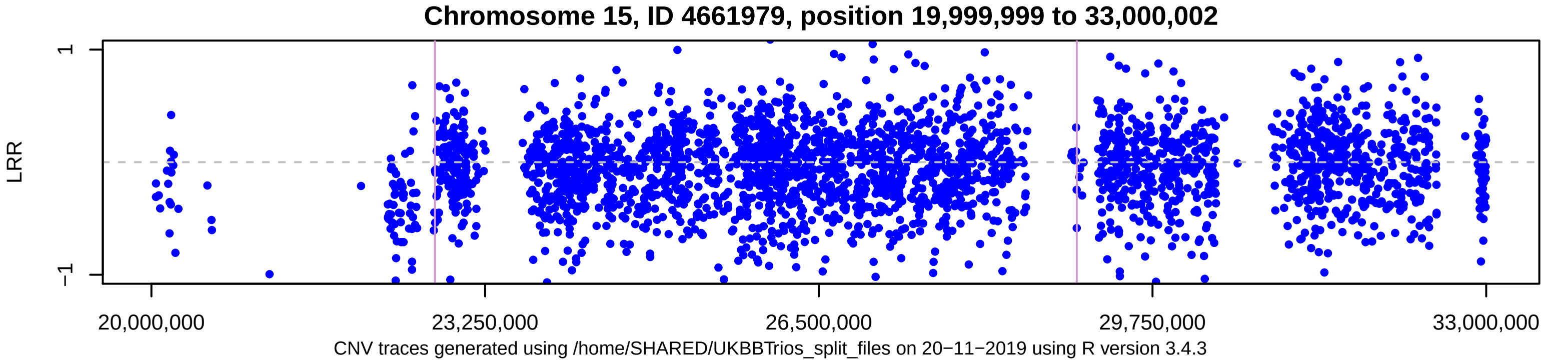


CNV traces generated using /home/SHARED/UKBBTrios_split_files on 20−11−2019 using R version 3.4.3

Chromosome 4, ID 2607929, position 100,033,629 to 104,161,402

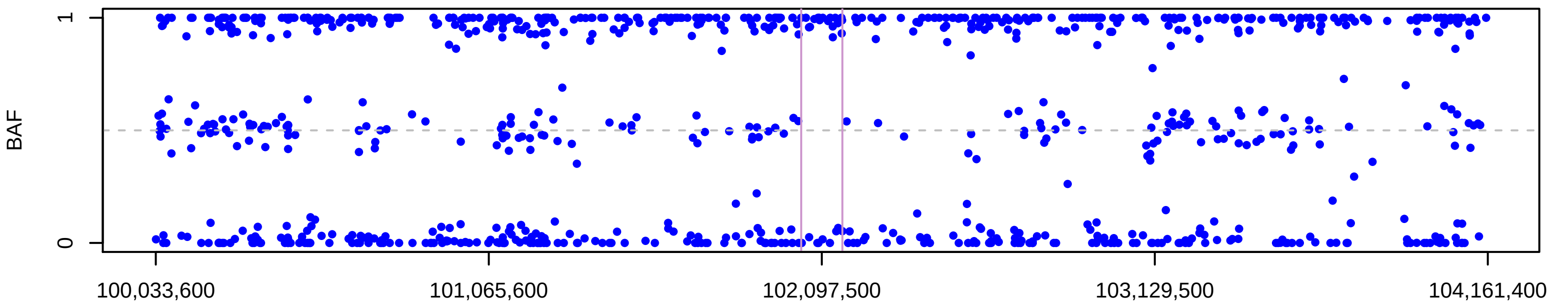


Chromosome 4, ID 2607929, position 100,033,629 to 104,161,402

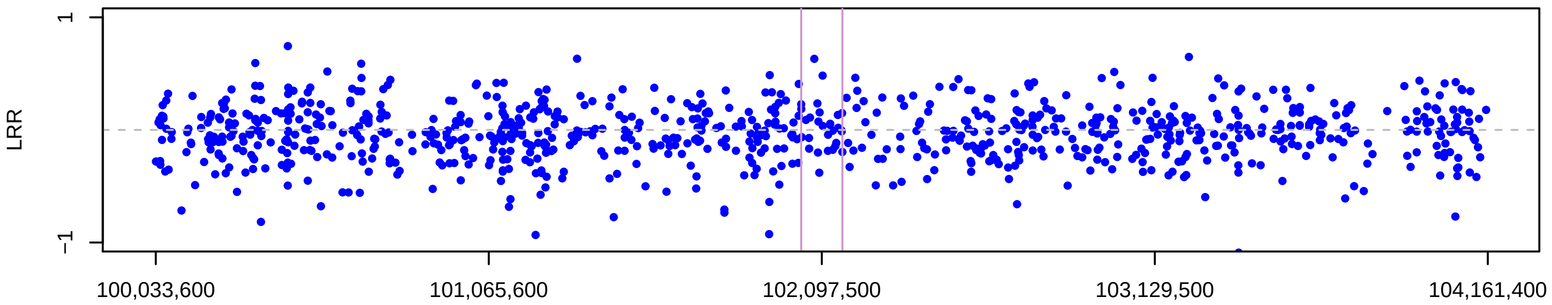


Chromosome 4, ID 1754684, position 100,033,629 to 104,161,402

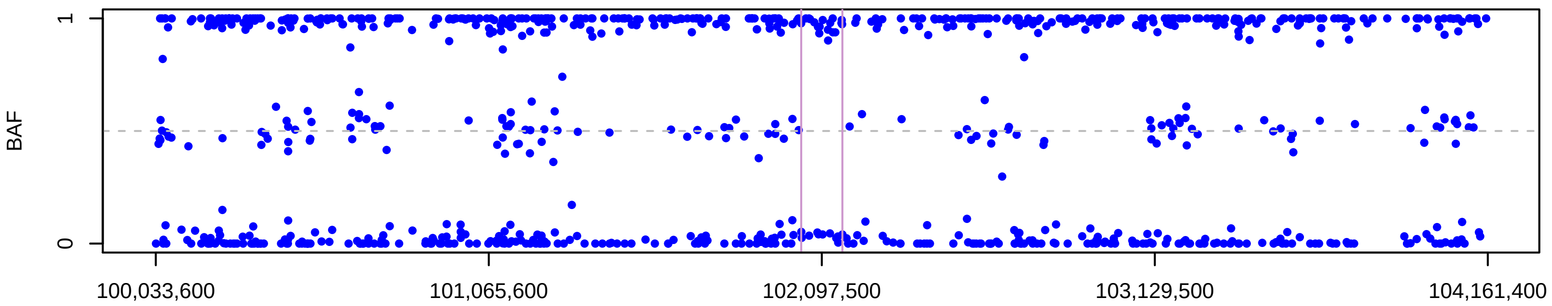


Chromosome 4, ID 1754684, position 100,033,629 to 104,161,402

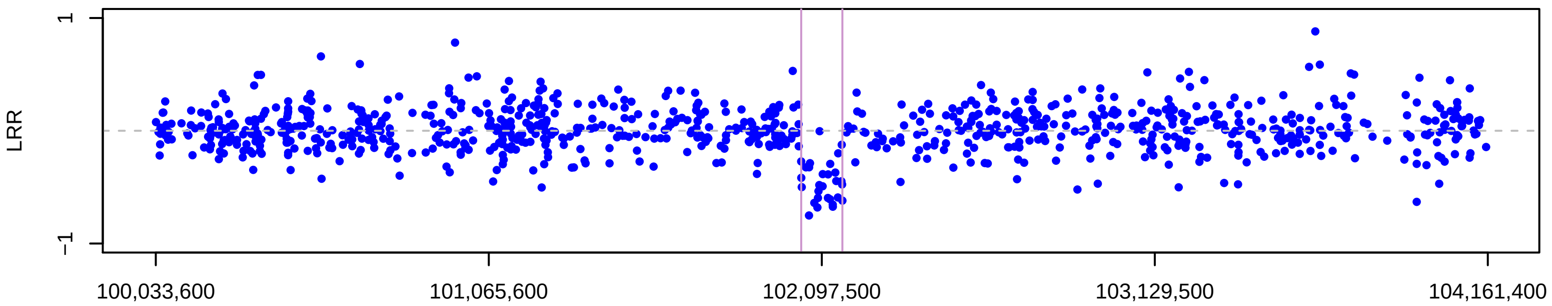


Chromosome 4, ID 1906870, position 100,033,629 to 104,161,402

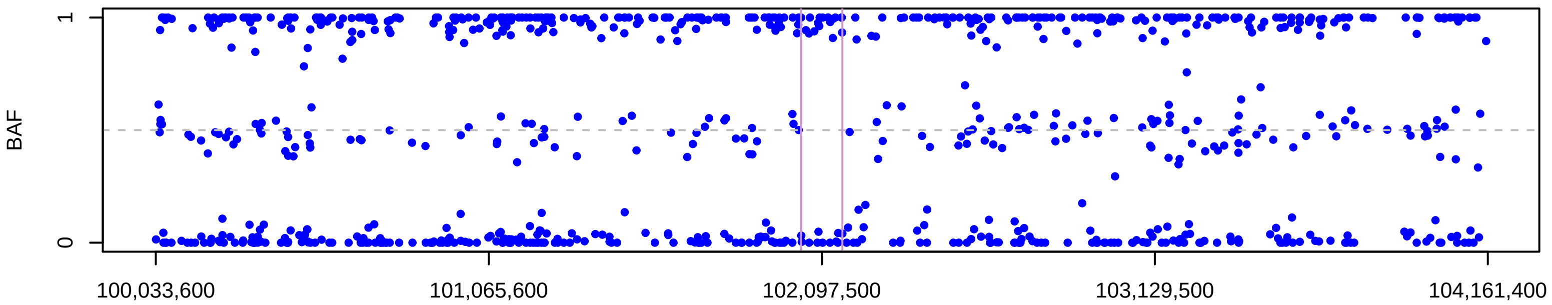


Chromosome 4, ID 1906870, position 100,033,629 to 104,161,402

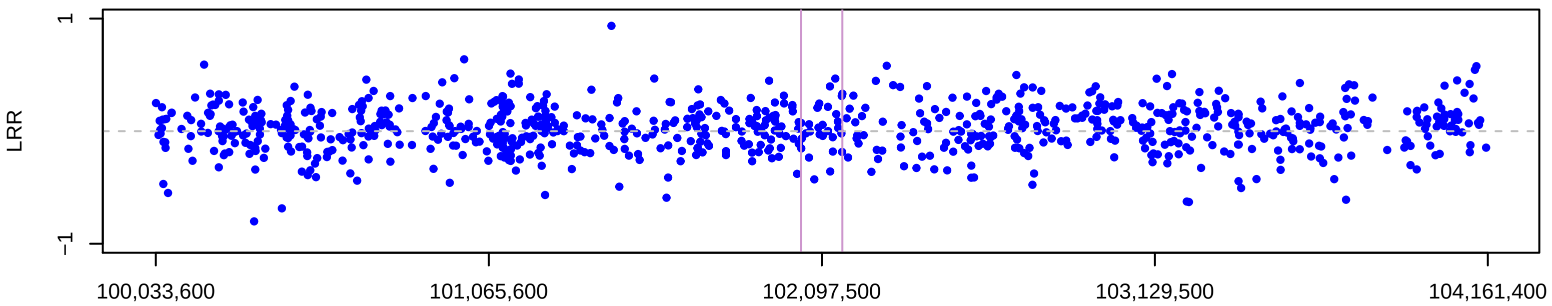


CNV traces generated using /home/SHARED/UKBBTrios_split_files on 20−11−2019 using R version 3.4.3

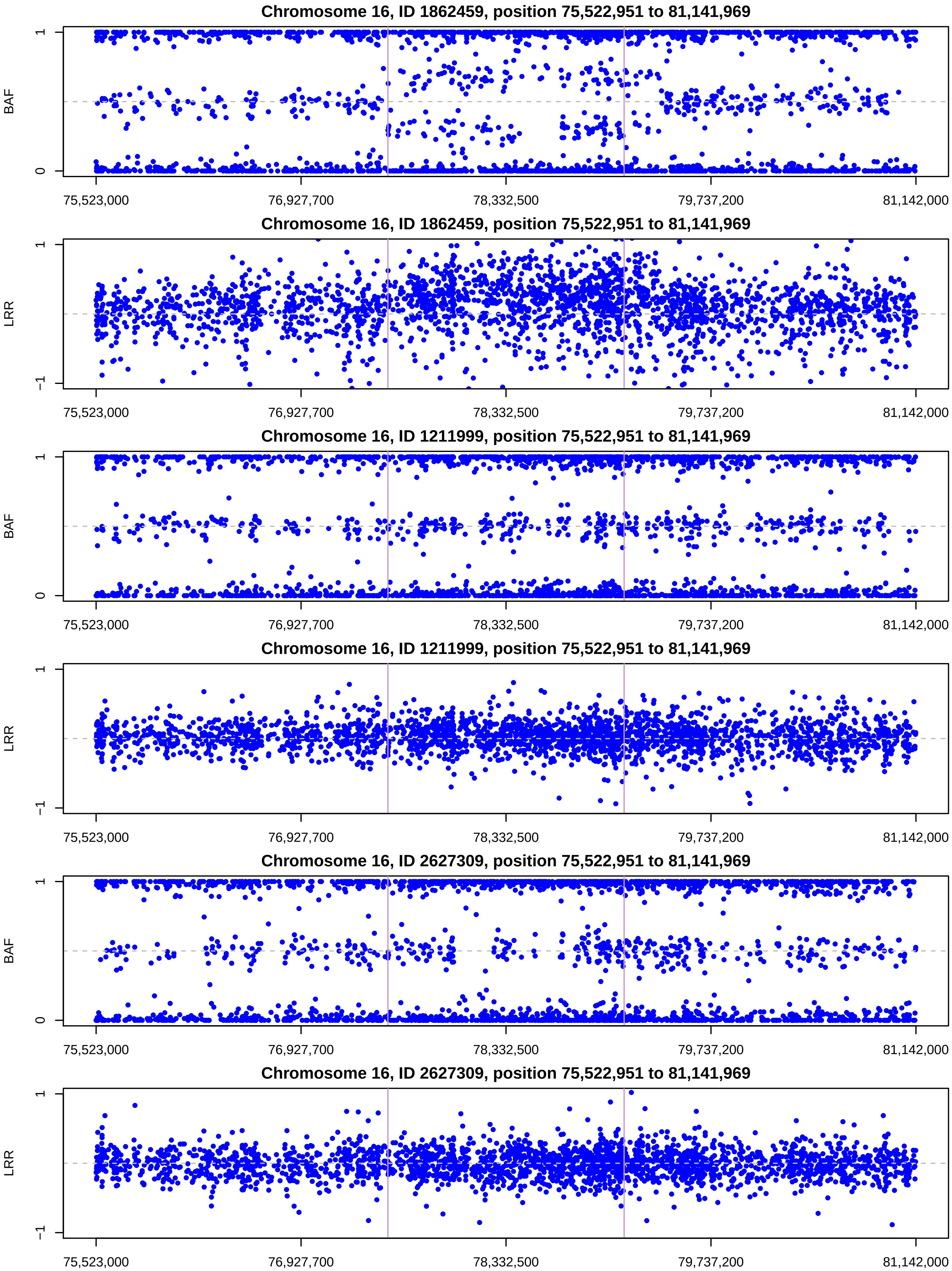


CNV traces generated using /home/SHARED/UKBBTrios_split_files on 03−05−2019 using R version 3.4.3

**Chromosome 19, ID 3298043, position 41,903,798 to 46,491,985**

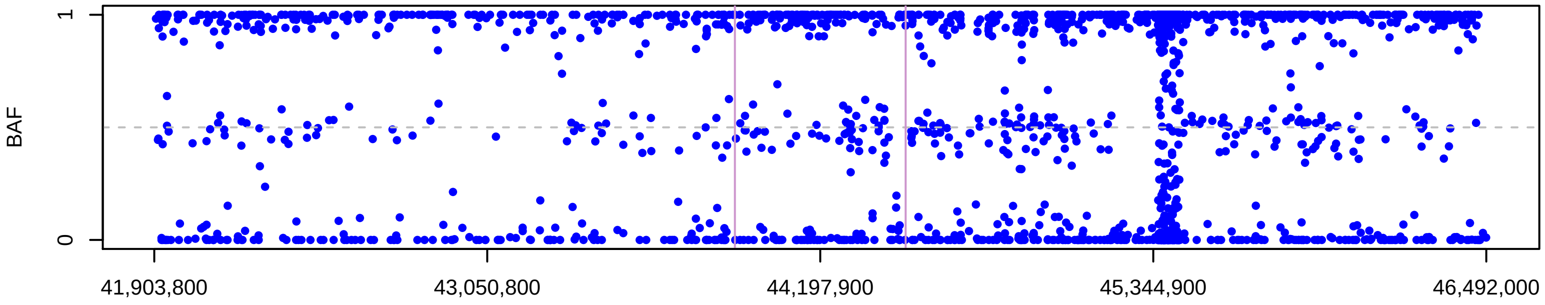


**Chromosome 19, ID 3298043, position 41,903,798 to 46,491,985**

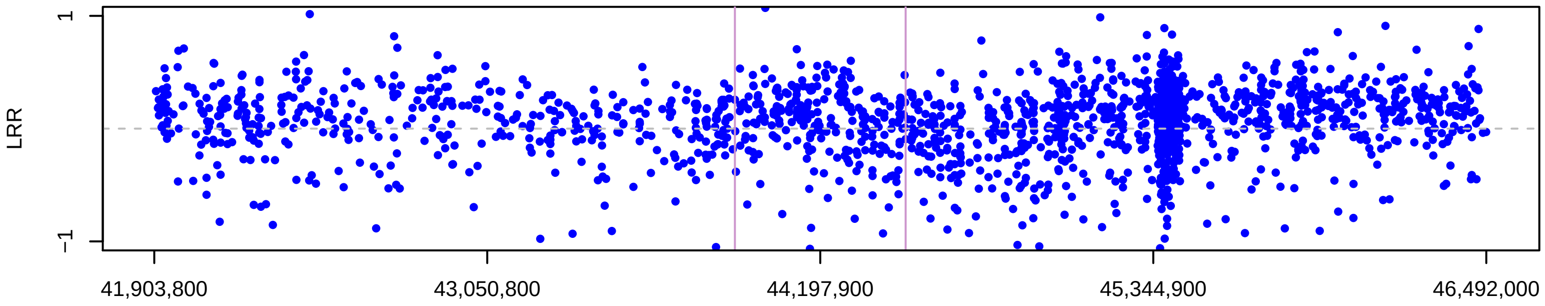


**Chromosome 19, ID 2883799, position 41,903,798 to 46,491,985**

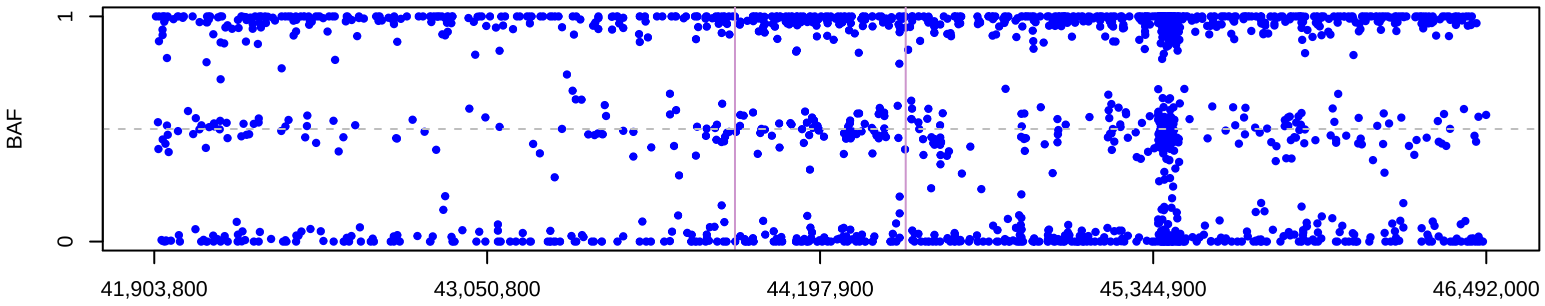


**Chromosome 19, ID 2883799, position 41,903,798 to 46,491,985**

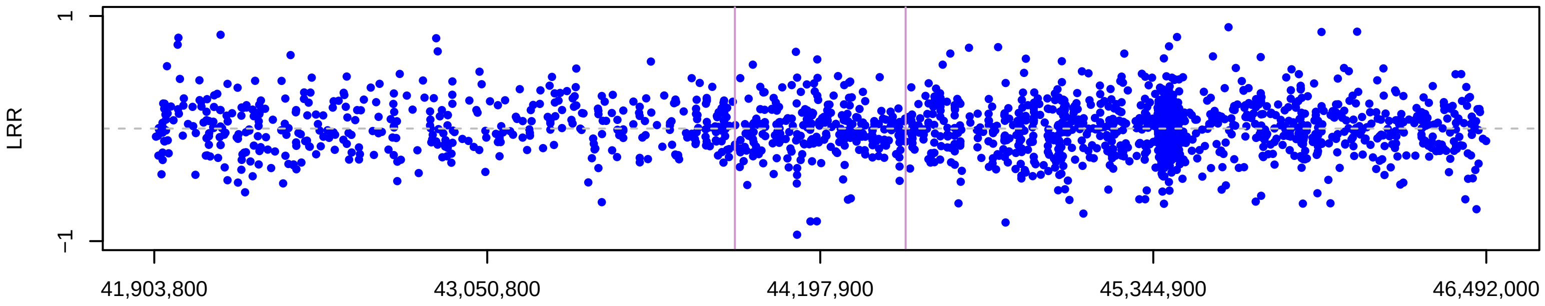


**Chromosome 19, ID 2095321, position 41,903,798 to 46,491,985**

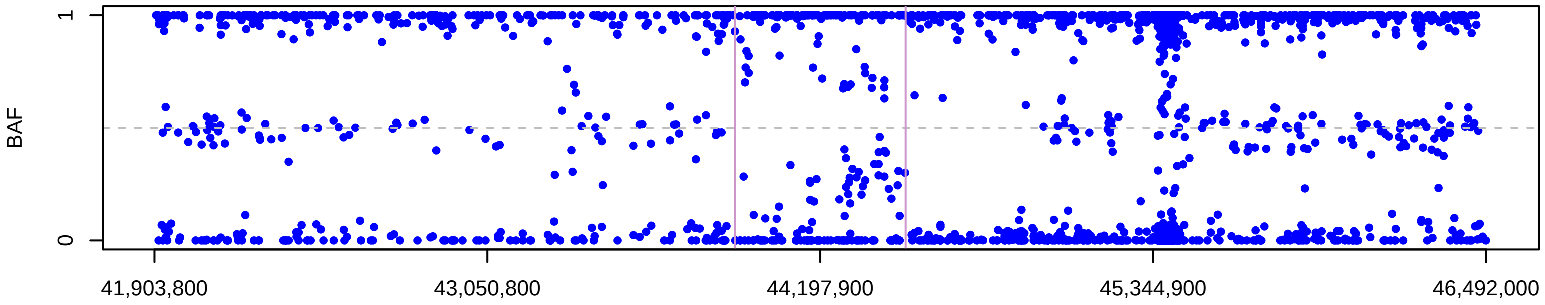


**Chromosome 19, ID 2095321, position 41,903,798 to 46,491,985**

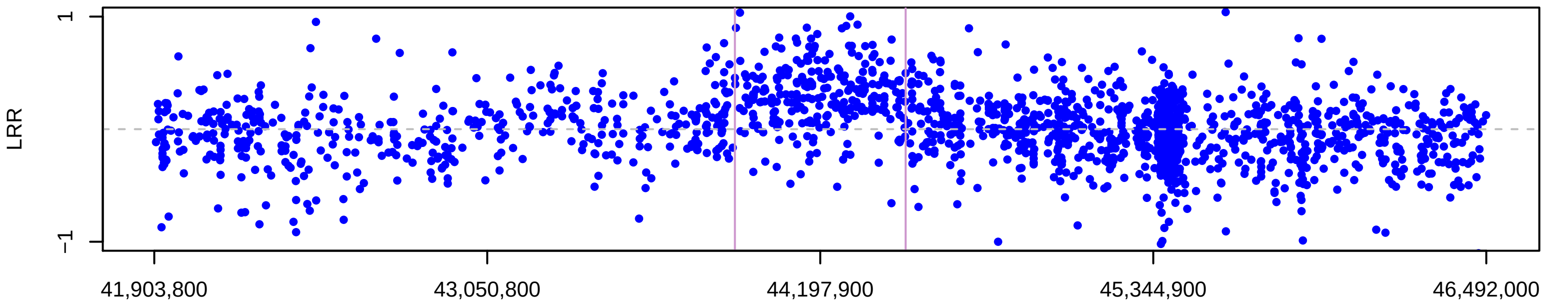


CNV traces generated using /home/SHARED/UKBBTrios_split_files on 20−11−2019 using R version 3.4.3

Chromosome 22, ID 2444969, position 16,876,483 to 23,505,589

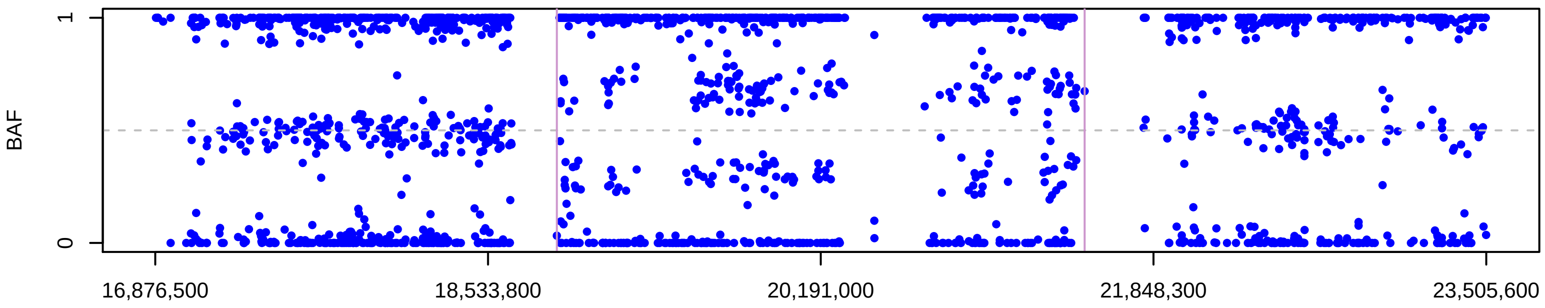


Chromosome 22, ID 2444969, position 16,876,483 to 23,505,589

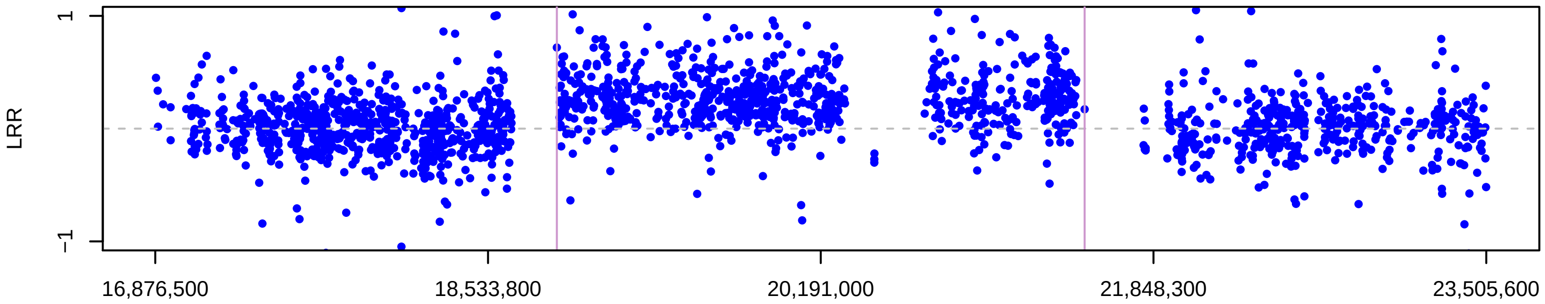


Chromosome 22, ID 4507686, position 16,876,483 to 23,505,589

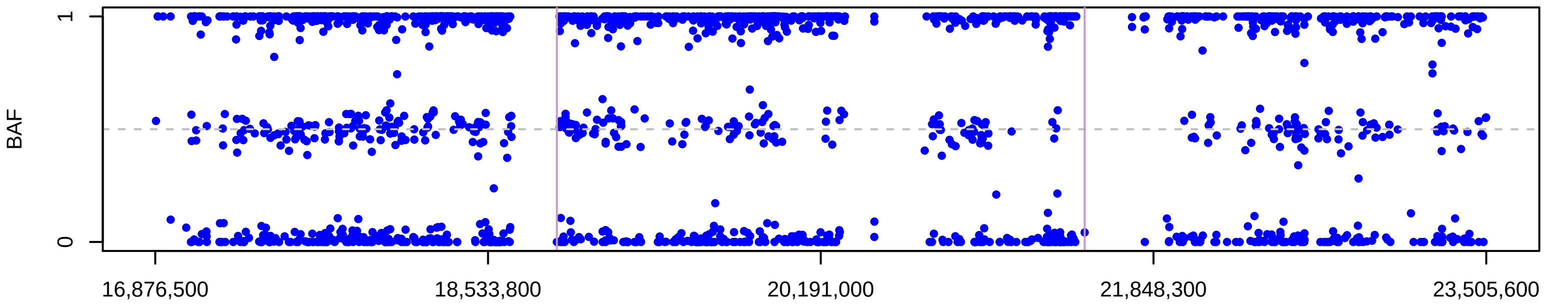


Chromosome 22, ID 4507686, position 16,876,483 to 23,505,589

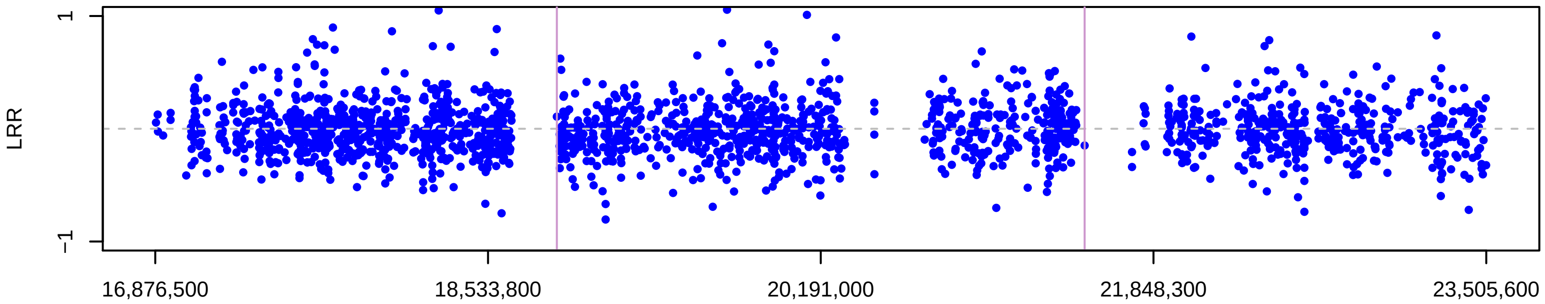


Chromosome 22, ID 2914764, position 16,876,483 to 23,505,589

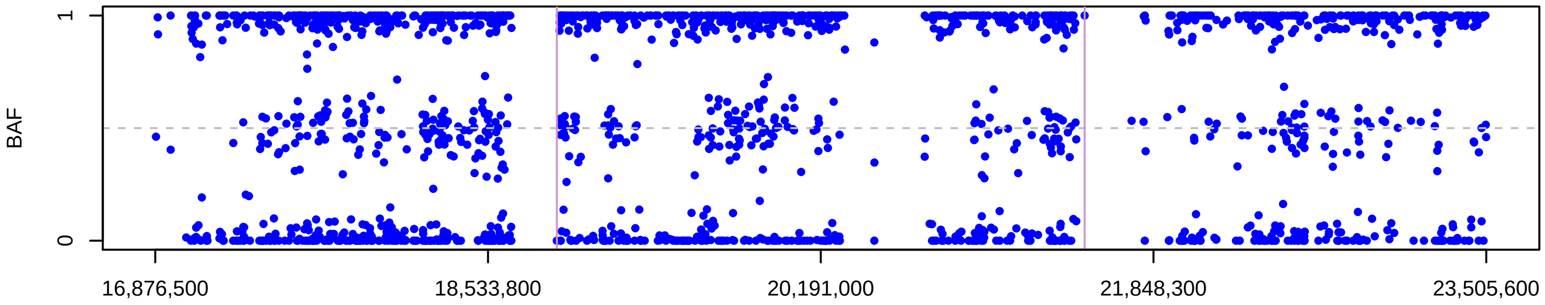


Chromosome 22, ID 2914764, position 16,876,483 to 23,505,589

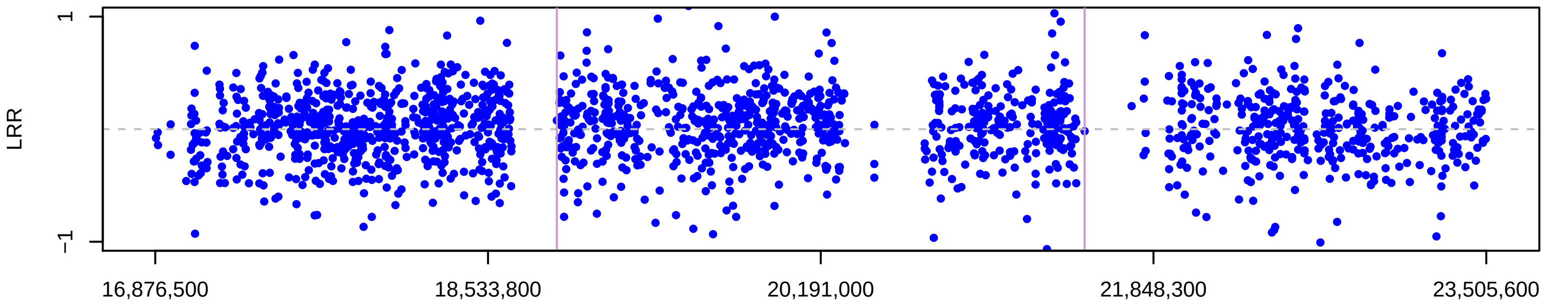


CNV traces generated using /home/SHARED/UKBBTrios_split_files on 20−11−2019 using R version 3.4.3

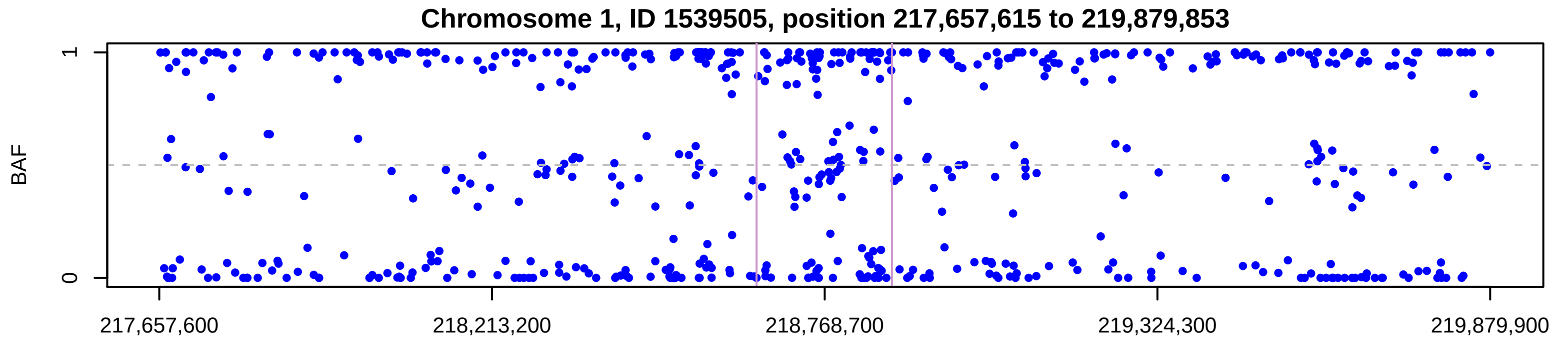


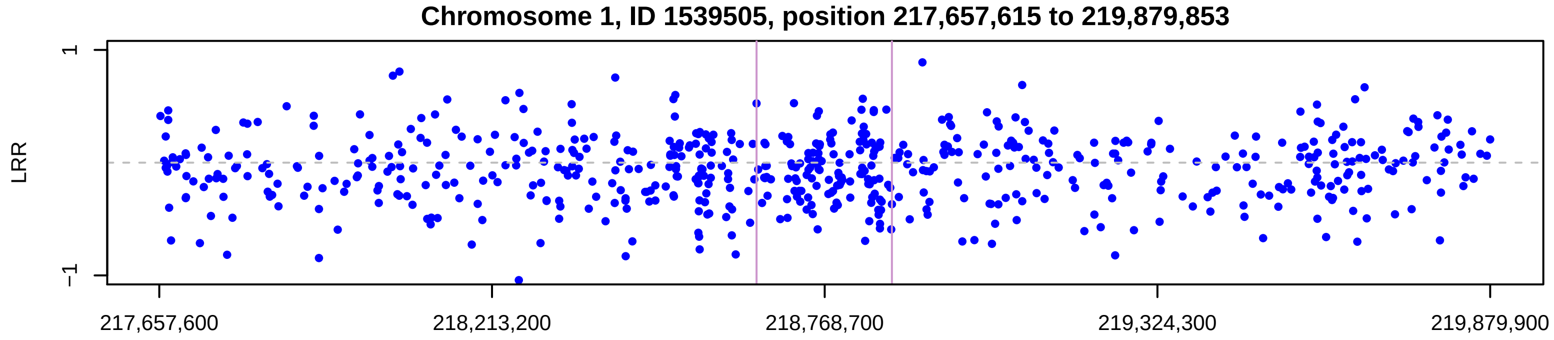


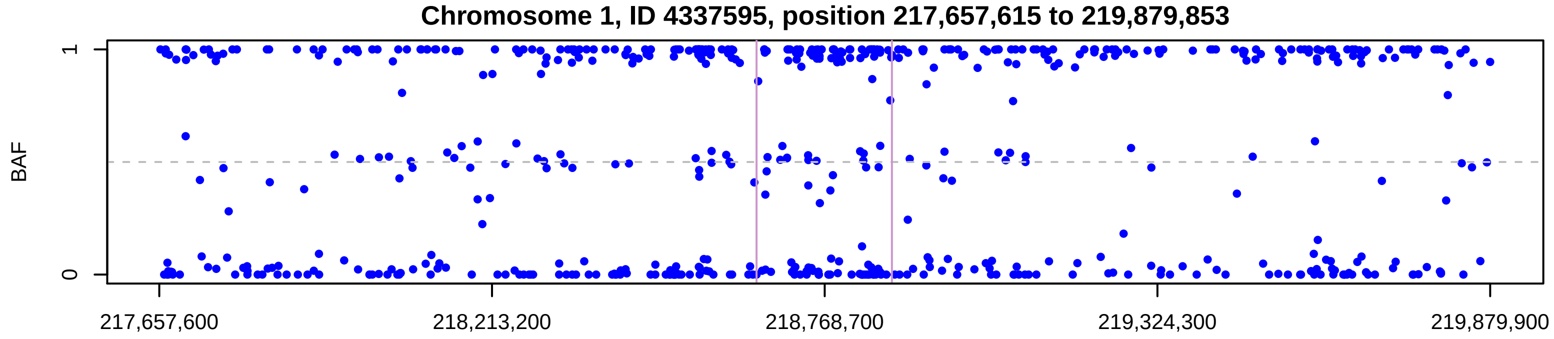


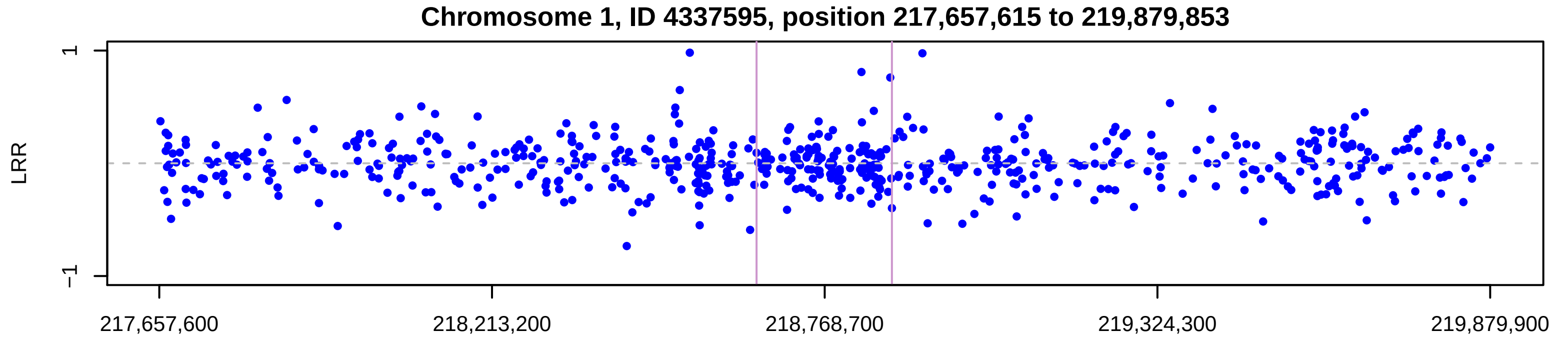


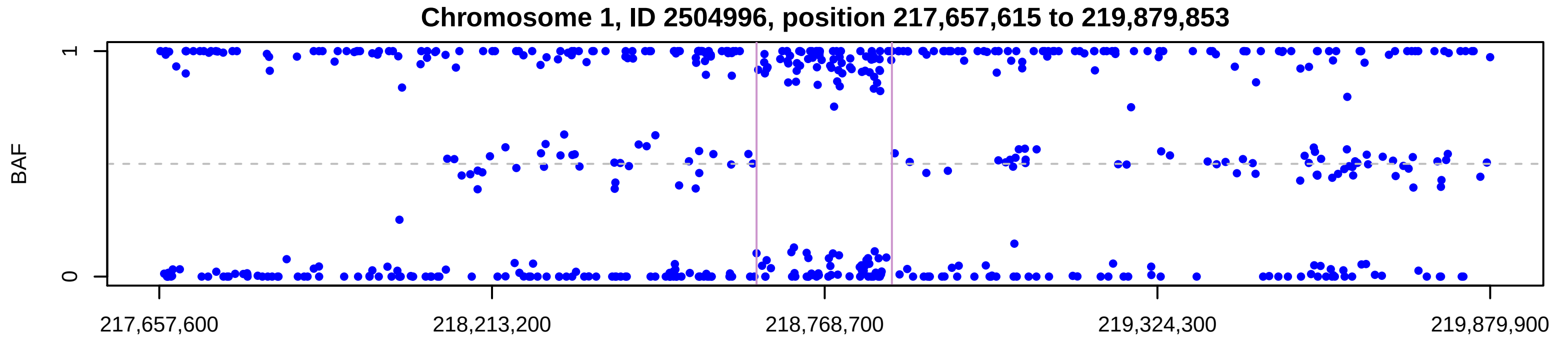


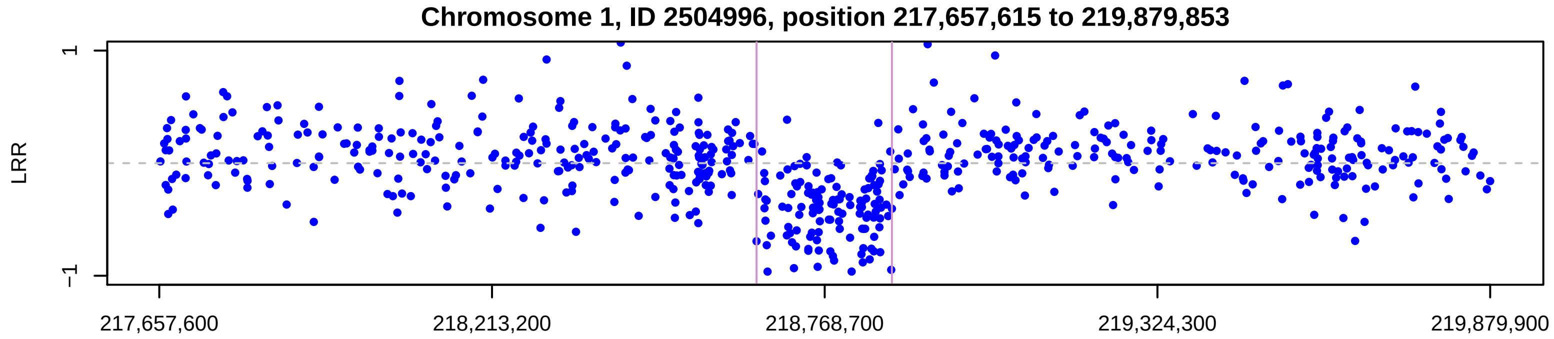


CNV traces generated using /home/SHARED/UKBBTrios_split_files on 07−05−2019 using R version 3.4.3

Chromosome 17, ID 2921432, position 32,816,054 to 38,249,386

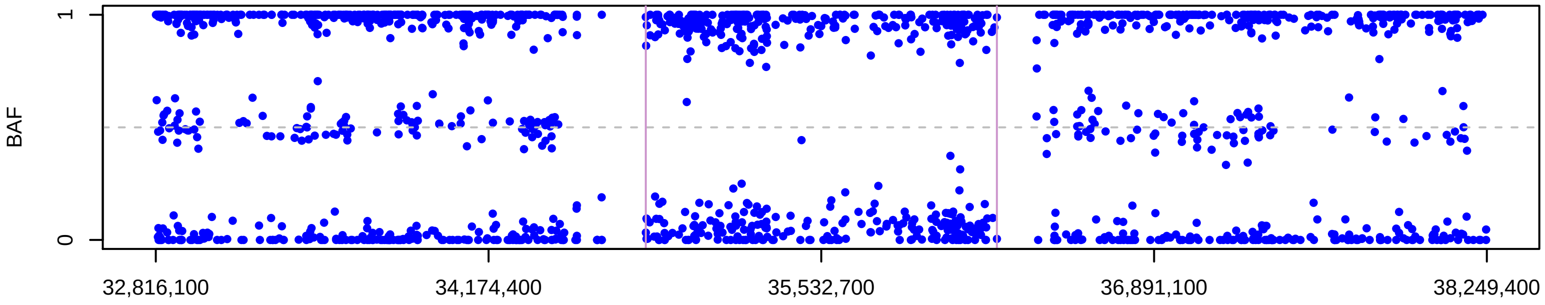


Chromosome 17, ID 2921432, position 32,816,054 to 38,249,386

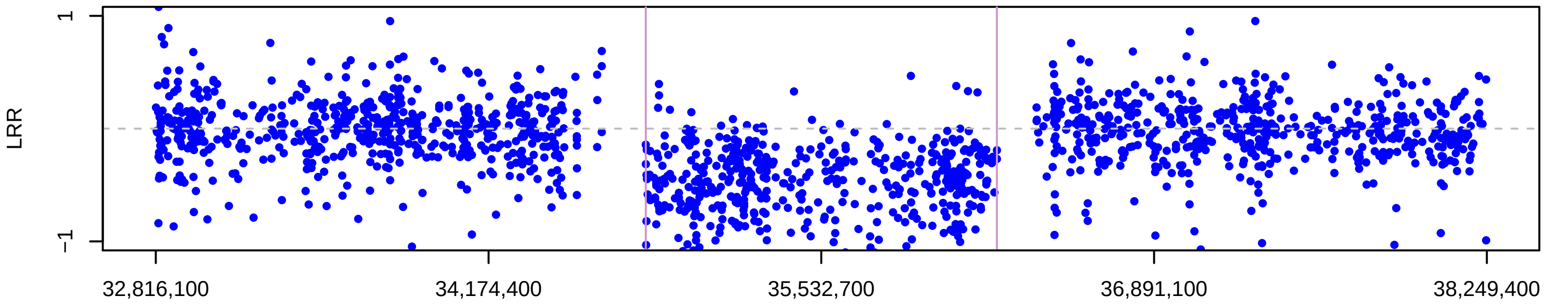


Chromosome 17, ID 4058117, position 32,816,054 to 38,249,386

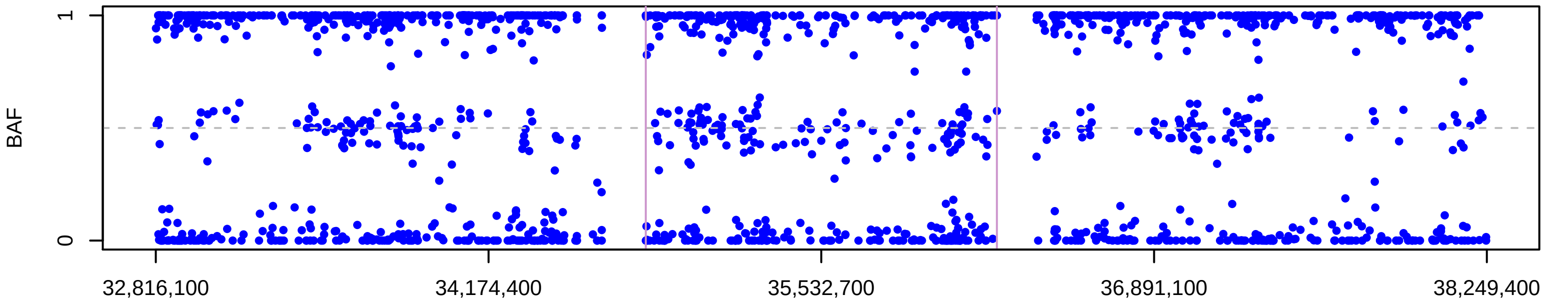


Chromosome 17, ID 4058117, position 32,816,054 to 38,249,386

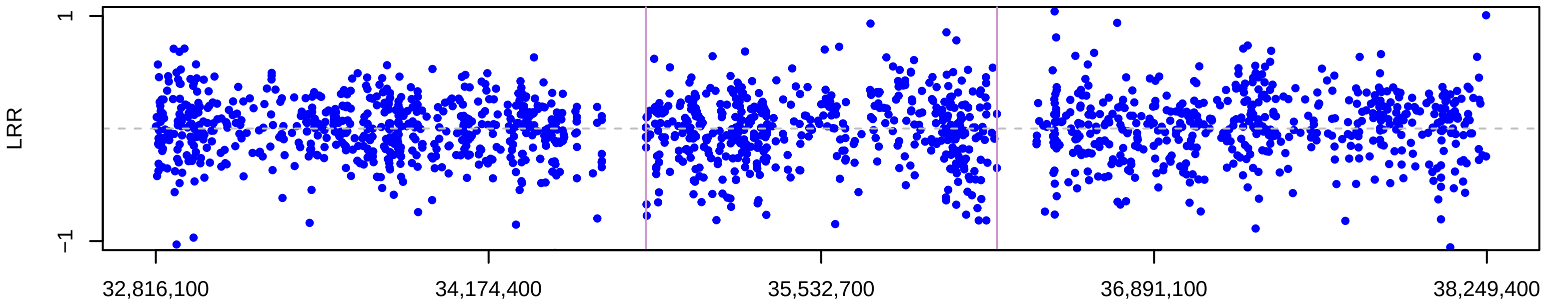


Chromosome 17, ID 2018806, position 32,816,054 to 38,249,386

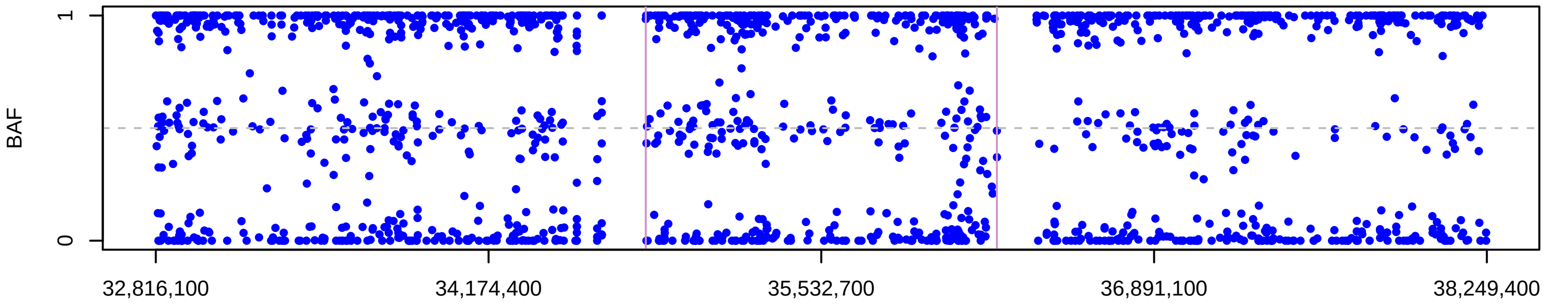


Chromosome 17, ID 2018806, position 32,816,054 to 38,249,386

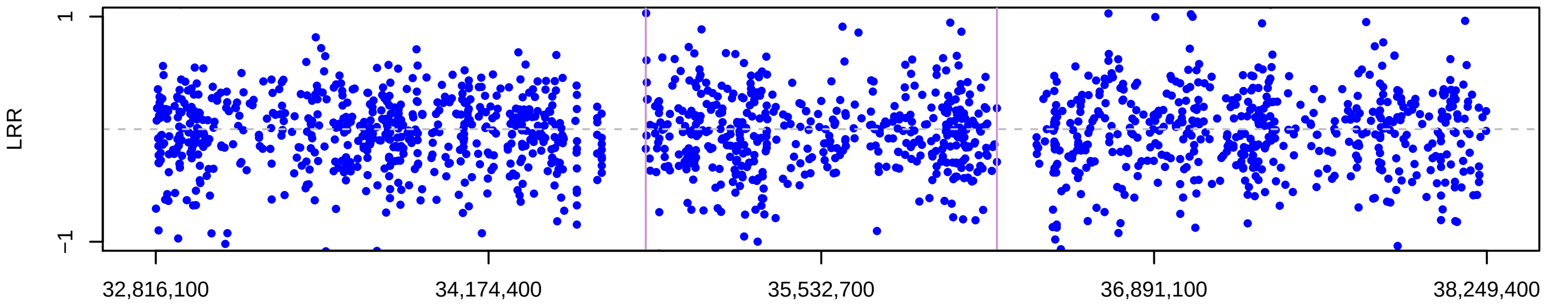


CNV traces generated using /home/SHARED/UKBBTrios_split_files on 20−11−2019 using R version 3.4.3

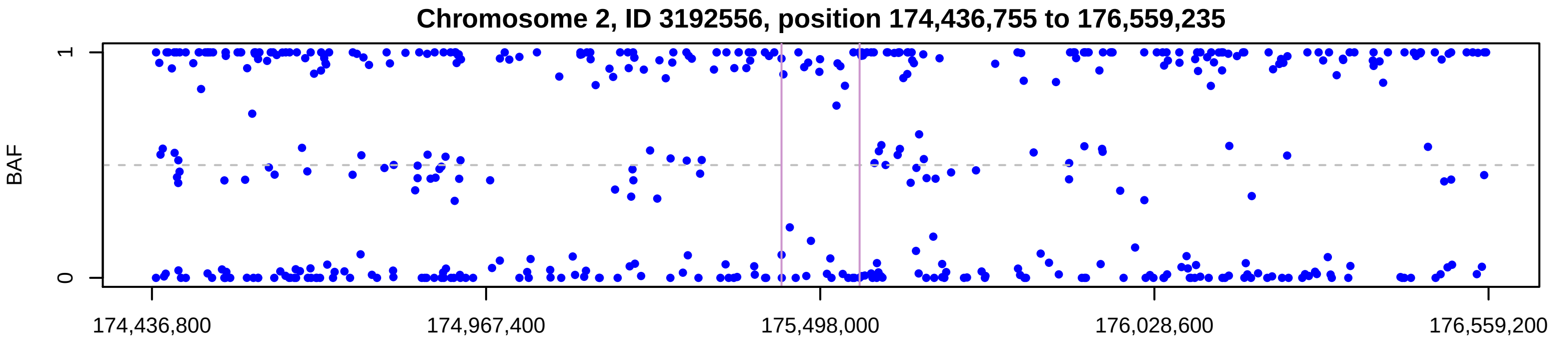


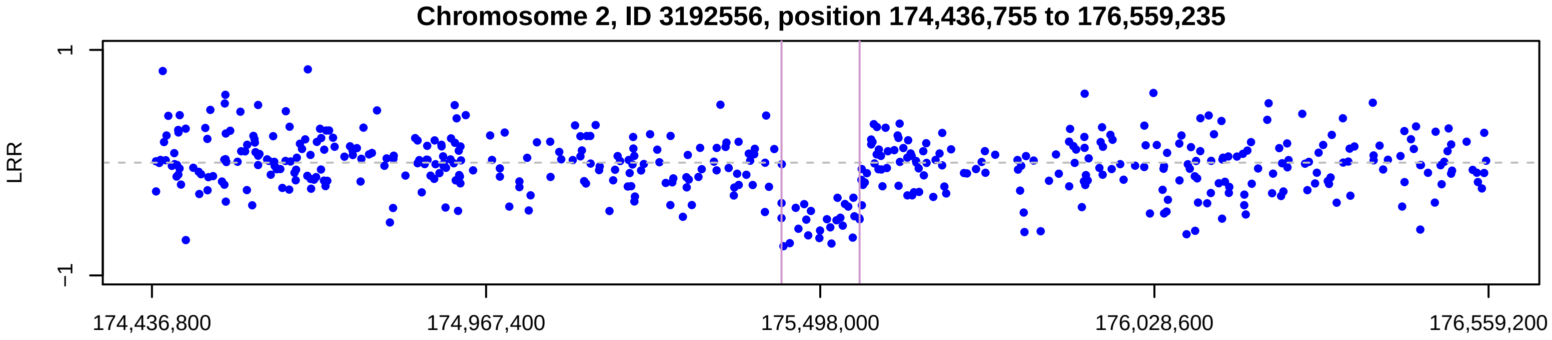


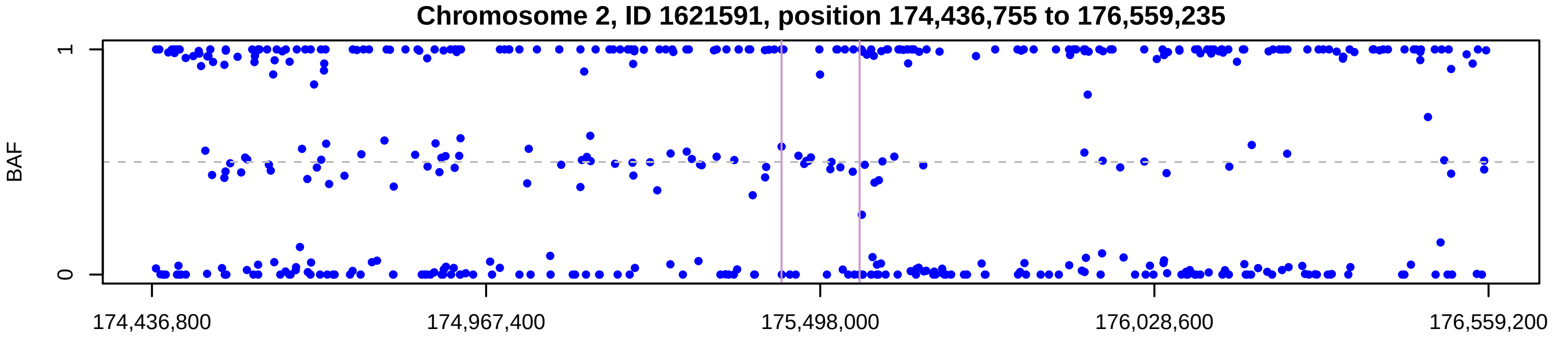


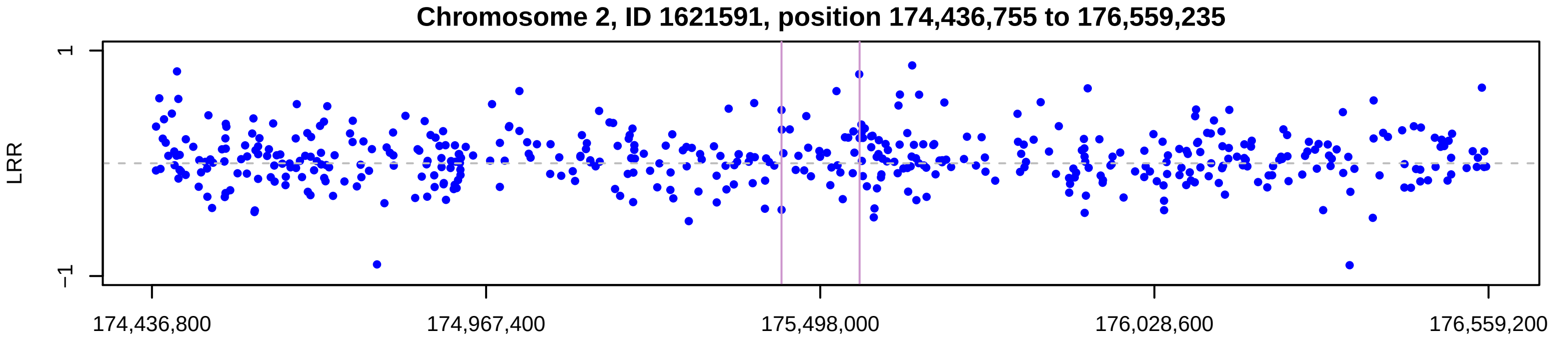


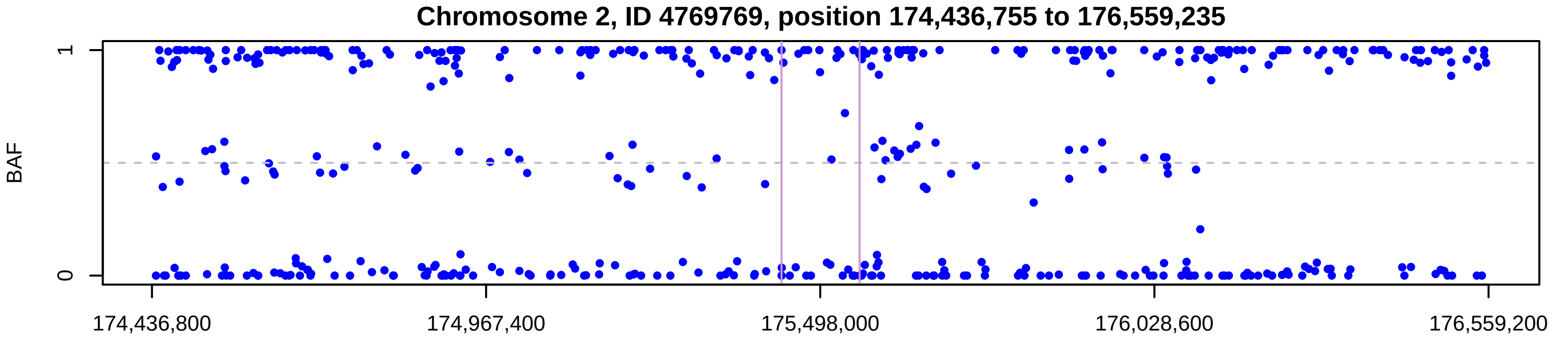


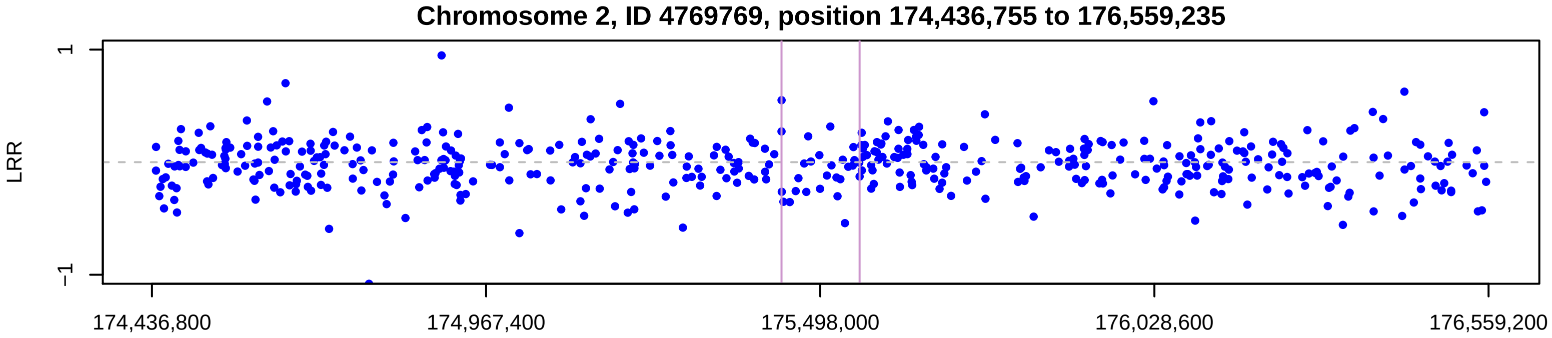


CNV traces generated using /home/SHARED/UKBBTrios_split_files/ on 13−05−2019 using R version 3.4.3

Chromosome 12, ID 2918552, position 7,796,656 to 8,323,309

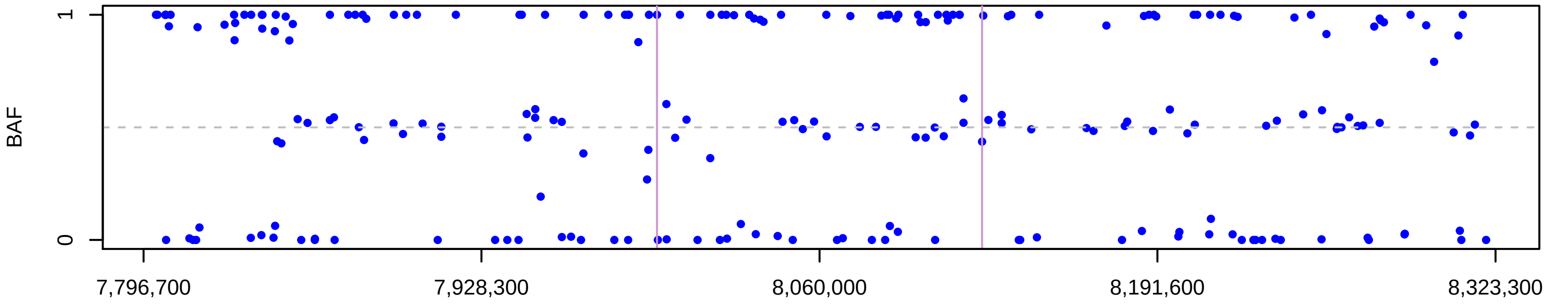


Chromosome 12, ID 2918552, position 7,796,656 to 8,323,309

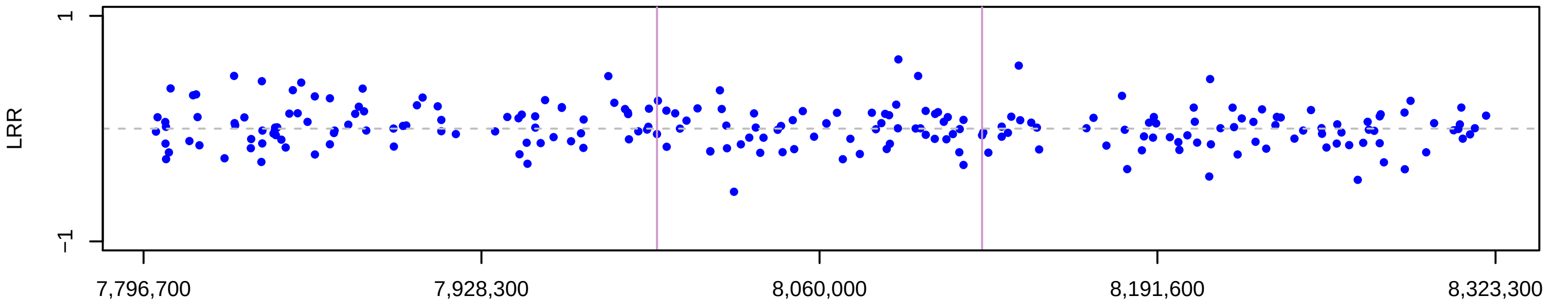


Chromosome 12, ID 4352466, position 7,796,587 to 8,323,309

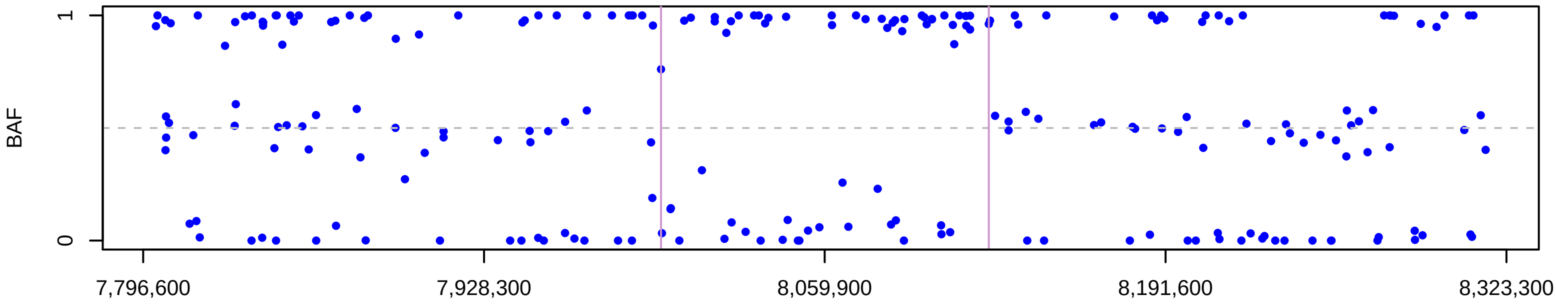


Chromosome 12, ID 4352466, position 7,796,587 to 8,323,309

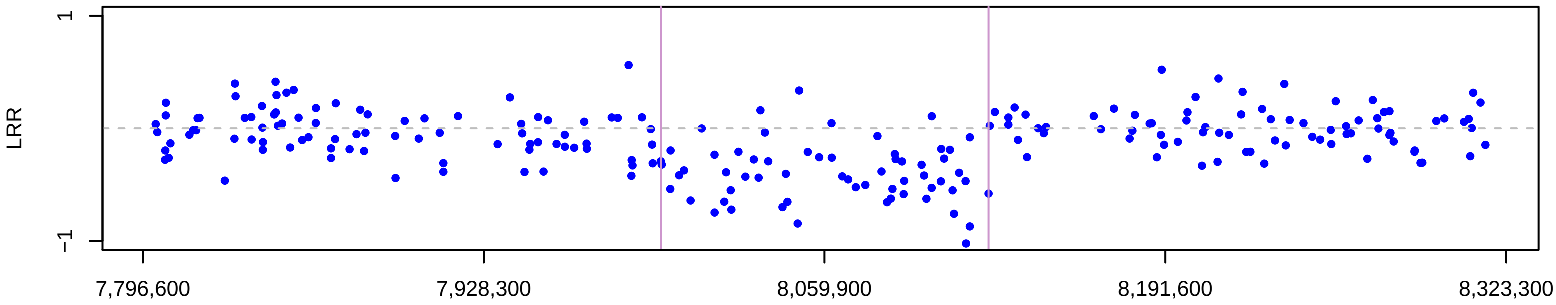


Chromosome 12, ID 4390290, position 7,796,587 to 8,323,309

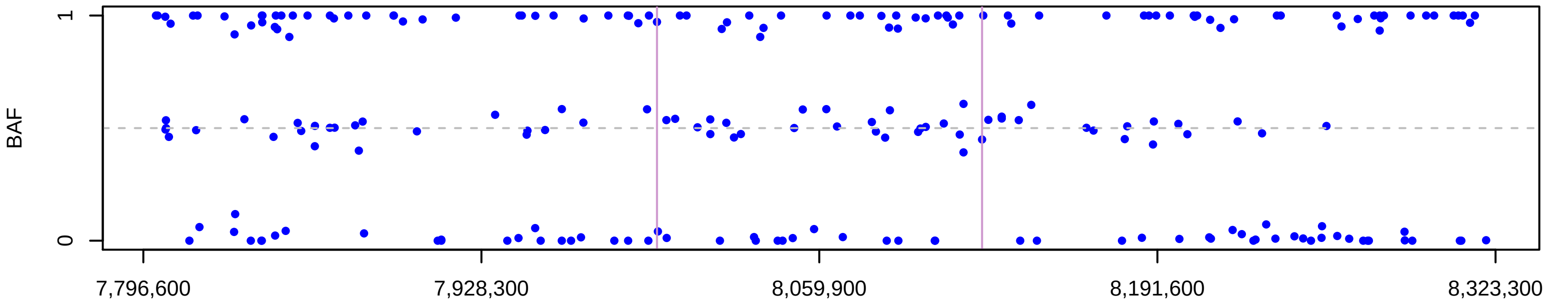


Chromosome 12, ID 4390290, position 7,796,587 to 8,323,309

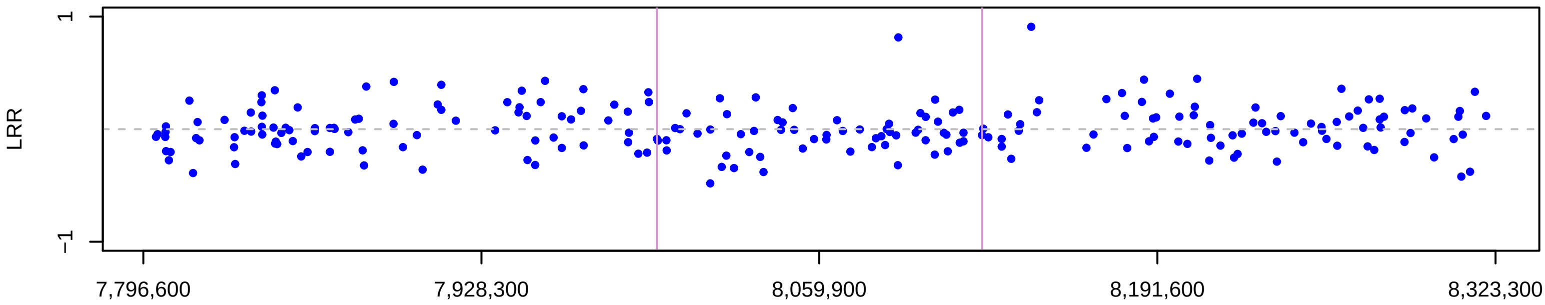


CNV traces generated using /home/SHARED/UKBBTrios_split_files on 20−11−2019 using R version 3.4.3

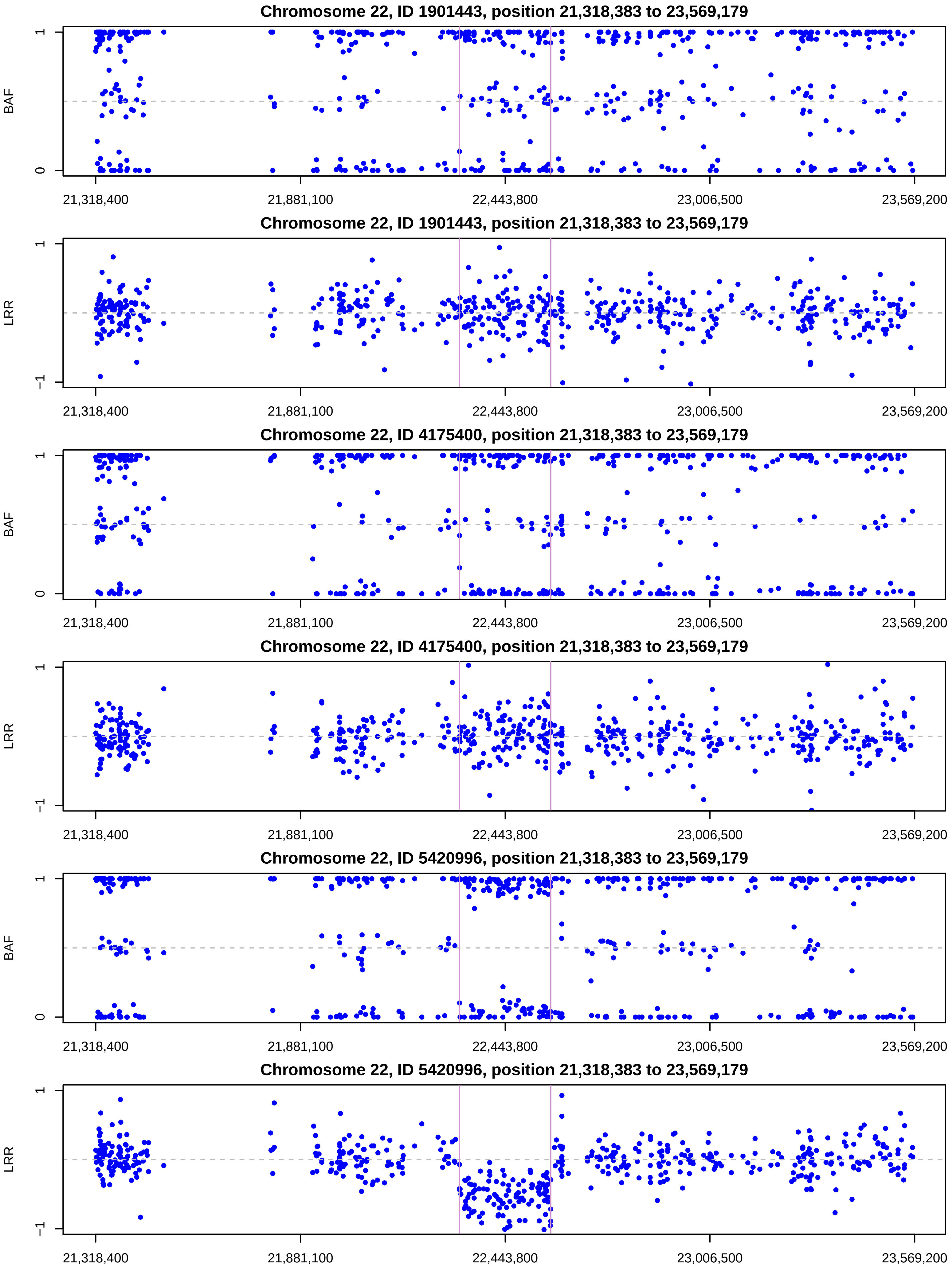

CNV traces generated using /home/SHARED/UKBBTrios_split_files/ on 13−05−2019 using R version 3.4.3
